# Siglec-engaging immunosuppressive sialoglycans are upregulated in prostate cancer and are targetable to suppress bone metastasis

**DOI:** 10.1101/2025.11.12.687981

**Authors:** Ziqian Peng, Kirsty Hodgson, Matthew Fisher, Shenglin Mei, Margarita Orozco-Moreno, Lizhi Cao, Michael Kemp, Wayne Gatlin, Louisa Donald, Feier Zeng, Michelle A Lawson, Daniel Ungar, David B Sykes, David J. Elliott, Li Peng, Ben Schumann, Ning Wang, Jennifer Munkley

**Author notes:** Correspondence to: Jennifer Munkley.

## Abstract

Prostate cancer is a leading cause of male cancer-related deaths over the age of 50. New treatment options for prostate cancer are urgently needed, especially for tumours that have spread to bone. Aberrant sialylation holds substantial potential for the discovery of new therapeutic targets but has remained relatively unexplored in the context of prostate cancer, primarily due to the lack of reliable reagents for detecting tumour sialoglycans in clinical tissue. Here, we address this knowledge gap using high-affinity Siglec-based sialoglycan-binding reagents (HYDRAs) to quantify tumour sialoglycans in tissues representing the full clinical heterogeneity of prostate tumours. Using HYDRA immunohistochemistry, we show that sialoglycans that can engage Siglec-3, -7, and -9 are upregulated in primary prostate cancer tissue and sialoglycan ligands for Siglec-7 correlate with prostate cancer bone metastasis and poorer patient prognosis. Analysis of prostate-derived tumours growing in bone reveals Siglec receptors are expressed by immune cells in the bone metastatic tumour microenvironment, suggesting that this axis may play a role in immune cell functions in bone metastatic prostate cancer. Indicating this is clinically actionable, an engineered bisialidase (E-612) can effectively strip Siglec ligands from prostate cancer cells and prolong survival times of mice with bone metastasis. Our findings identify a novel mechanism involving Siglec-engaging sialoglycans in driving the growth of prostate cancer bone metastasis and demonstrate how this axis can be targeted to impede lethal prostate cancer progression.

## Introduction

Prostate cancer is a common cancer in men leading to more than 350,000 deaths worldwide every year [1]. When prostate cancer is localised, curative therapies are available and the 5-year survival rate is 97%, but for patients with metastasis, 5-year survival can be as low as 32% [2]. The growth of prostate tumours is driven by androgen receptor (AR) signalling, and initial therapeutic options for advanced prostate cancer are hormone-based therapies such as anti-androgens [3–5]. However, eventually most tumours will become resistant to hormone therapy and progress to lethal castrate resistant prostate cancer (CRPC) [6]. Bone metastasis is common in prostate cancer, with up to 80% of patients with advanced disease diagnosed with tumours that have spread to the skeleton [2]. Although it is possible to manage the symptoms [7], no curative treatments are available for prostate cancer that has spread to bone and there is a critical need to develop new therapeutic options.

An area of innovation in the search for new cancer therapies is the targeting of cancer-associated glycosylation [8, 9]. Glycans decorate all cells and most secreted proteins and play essential roles in all biological processes [10]. Cancer cells have a different ‘glycan coat’ than healthy cells [11, 12], and this aberrant glycosylation plays a role in all of the cancer hallmarks [12]. A common feature of tumours is an increased density of glycan structures terminating in sialic acid (sialoglycans), including truncated *O*-glycans, as well as an overexpression of tri- and tetra-antennary *N*-glycans [13, 14]. Altered sialylation can promote tumour cell survival and metastasis, immune evasion, and the promotion of drug resistance [13, 15, 16]. Sialoglycans can inhibit apoptosis by modifying cell surface receptors, including Fas and TNFR1, through preventing the formation of death-inducing signalling complexes [17, 18]. Aberrant sialylation also contributes to metastasis by promoting cell detachment and driving the spread of malignant cells, for example by modulating EGFR signalling [19] or the EMT pathway [20]. Sialoglycan-covered antigens can also act as ligands for receptors called Siglecs (sialic acid–binding immunoglobulin-like lectins) that are expressed on immune cells, leading to immunosuppressive signalling that promotes immune evasion [16]. Humans express 14 unique Siglecs that have distinct preferred sialoglycan ligands [21]. The role of Siglecs in cancer appears to be diverse and complex, with the tumour glyco-code consisting of a heterogenous mixture of sialoglycans binding to Siglecs via both low affinity and high avidity interactions [22]. Since sialoglycans can fuel cancer progression, they are attractive targets for novel cancer therapies. Strategies to block aberrant sialylation in tumours, including sialylation inhibitors, Siglec blocking antibodies, and targeted sialidases, are being actively pursued [13].

The prostate cancer glycome, which includes all glycans synthesised by cells, has remained relatively understudied. However, glycans hold great potential as biomarkers and therapeutic targets [23, 24]. Studies have functionally linked altered sialylation to prostate cancer growth, bone metastasis and anti-tumour immunity [25–27], and provide proof of principle data demonstrating the potential to target sialylated glycans to impede prostate cancer progression [26, 28]. ST6GAL1-mediated α2,6 sialylation of *N*-glycans can drive prostate cancer growth and promote the spread of tumours to bone [25, 26], and ST3GAL1, which adds sialic acid residues in α2,3 linkage to *O*-glycans, can synthesise Siglec ligands and mediate anti-tumour immunity in prostate cancer [27]. *N*-glycan mass spectrometry imaging [29] has identified significant changes to the glycosylation patterns in various stages of prostate cancer, including alterations to sialylated *N*-glycans in advanced disease [25, 26, 30, 31]. Emerging research suggests that sialoglycans with the capacity to engage Siglec receptors may be upregulated on the surface of prostate cancer cells [22, 27, 32]. However, a correlation between sialoglycans that can engage Siglec receptors and prostate cancer progression has not yet been reported. Using Siglecs to reliably assess Siglec-engaging immunosuppressive sialoglycans is challenging due to the low Siglec to ligand binding affinity [21]. Therefore, comprehensive studies are needed that reliably monitor the expression of these ligands across the full heterogeneity of clinical prostate cancer tissue. Previous studies show that targeting the ligands for Siglec-7/9 (by knockout of the glycosyltransferase ST3GAL1 or by using Siglec-7/9 blocking antibodies) can decrease prostate tumour growth and enhance anti-tumour immunity [27, 32]. We have previously shown that pre-treatment of prostate cancer cells with a sialyltransferase inhibitor removes sialylated glycans and can suppress bone metastasis [26]. These findings raise the possibility that systemic therapies to disrupt the Siglec-sialoglycan axis will likely be effective against bone metastatic prostate cancer. However, this has not yet been evaluated using pre-clinical mouse models.

Here, using proprietary sialoglycan binding reagents, we monitored immunosuppressive sialoglycans in seven well annotated prostate cancer tissue microarrays (TMAs), allowing us to investigate Siglec sialoglycan ligand expression across the full clinical heterogeneity of clinical prostate tumour tissues. Sialoglycan binding reagents provided by Palleon Pharmaceuticals consist of multimeric Siglec domains to achieve superior avidity [33]. Our findings reveal a significant upregulation of immunosuppressive sialoglycans, including ligands for Siglec-3, -7, and -9, in primary prostate cancer tissues relative to matched control samples and in therapy resistant prostate tumours growing in bone. We show that sialoglycan ligands for Siglec-7 correlate with bone metastasis and reduced survival times in prostate cancer patients. We also reveal that Siglec receptors are expressed by immune cells in the bone metastatic prostate tumour immune microenvironment (TIME). Furthermore, we demonstrate that an engineered dual-action sialidase (E-612), that strips Siglec ligands from prostate cancer cells, prolongs survival times of mice with prostate cancer bone metastasis. Our study identifies a novel mechanism involving Siglec-engaging sialoglycans as driving the growth of bone metastatic prostate tumours and indicates the Siglec-sialoglycan axis can be clinically targeted to impede prostate cancer progression.

## Results

### 1. Sialoglycans that can engage Siglec-3, -7, and -9 are upregulated in prostate tumours

Previous studies suggest that sialoglycans with the capacity to engage Siglec-7 and Siglec-9 are upregulated on the surface of prostate cancer cells [22, 27, 32]. However, these findings were based on small sample sizes (<50 clinical tissues) and did not investigate correlations between sialoglycans that can engage Siglec receptors and prostate cancer progression.

The Siglec-Fc proteins utilised to detect sialoglycans, while valuable tools, have reported limitations for ligand specificity and avidity [34]. To overcome this, we utilised the novel HYDRA reagents (Palleon Pharmaceuticals) including HYDRA-3, -7, and -9 (that consist of multimeric Siglec domains with higher avidity) to detect sialoglycans recognised by Siglec-3, -7, and -9 respectively [22, 33]. Our study was focused on profiling Siglec-3, -7, and -9 sialoglycan ligand expression on prostate tumours, since among the 14 Siglecs in humans, these are the major inhibitory Siglecs on both innate and adaptive immune cells [22]. Using HYDRA-3, -7, and -9 immunohistochemistry, we monitored the levels of sialoglycans that can engage Siglec-3, -7, and -9 across three different tissue microarrays (TMAs). First, a TMA with tissue samples from 40 prostate cancer patients (TMA cohort 1) showed that all three sialoglycan types are expressed at significantly higher levels in prostate cancer tissue relative to normal prostate tissue (Figure 1A). Second, analysis of a 96 case TMA containing 17 normal prostate tissue samples and 79 samples of prostate tumour tissue (TMA cohort 2) showed that sialoglycan ligands for Siglec-3, -7, and -9 are significantly upregulated in prostate tumours (Figure 1B). Finally, staining of a TMA comprising matched normal and prostate cancer tissue from 200 patients (TMA cohort 3) showed that sialoglycan ligands for Siglec-3 (paired t test, p<0.0001), Siglec-7 (paired t test, p=0.0003) and Siglec-9 (p<0.0001) are found at significantly higher levels in prostate cancer tissue relative to matched normal tissue from the same patient (Figure 1C). Taken together, these novel molecular reagents reveal upregulation of sialoglycans that engage Siglec-3, -7, and -9 in clinical prostate cancer tissue.

**Figure 1.**
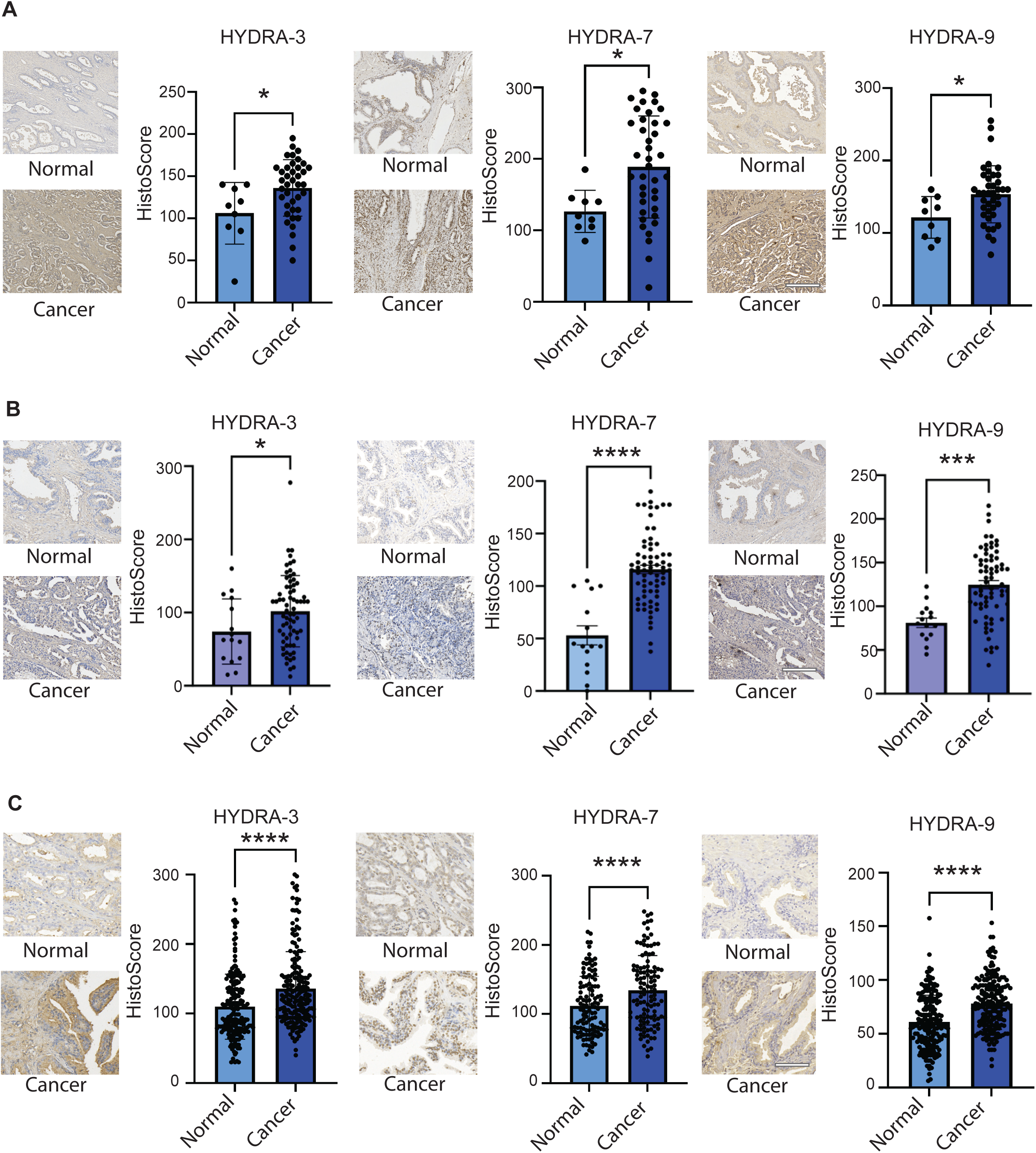
Prostate tumours have increased HYDRA scores compared to normal prostate tissue. (**A**) Analysis of Siglec-engaging sialoglycans using HYDRA immunohistochemistry in a TMA comprising 51 prostate tissue samples shows ligands for Siglec-3 (unpaired t test, p=0.0216), Siglec-7 (unpaired t test, p=0.0143) and Siglec-9 (unpaired t test, p=0.0271) are upregulated in prostate tumour tissue relative to normal prostate tissue. (**B**) Staining a previously published 96 case TMA [66, 72] containing 17 normal prostate tissue samples and 79 samples of prostate tumour tissue showed that sialoglycan ligands for Siglec-3 (unpaired t test, p<0.001), Siglec-7 (unpaired t test, p<0.0001) and Siglec-9 (unpaired t test, p0.0171) are found at significantly higher levels in prostate tumours compared to normal prostate tissues. (**C**) HYDRA immunohistochemistry analysis of Siglec-3, -7 and -9 ligands in a previously published 200 case TMA [66, 72] containing matched tumour and normal tissues from the same patient. Sialoglycan ligands recognised by Siglec-3 (paired t test, p<0.0001), Siglec-7 (paired t test, p=0.0003) and Siglec-9 (p<0.0001) are significantly increased in prostate cancer tissue relative to matched normal tissue from the same patient. Scale bar is 200□µm.

### 2. Expression of Siglec-7 engaging sialoglycans correlate with bone metastasis and reduced survival times in patients

We next monitored ligands for Siglec-3, -7, and -9 across five patient cohorts to investigate if immunosuppressive sialoglycans might also correlate with more aggressive prostate tumours. Analysis of the 96 case TMA cohort shown in Figure 1B revealed higher expression of ligands for Siglec-3, -7, and -9 in high grade (Gleason 4-5) tumours compared to lower grade (Gleason 1-3) tumours (Supplementary Figure 1). We also used HYDRA immunohistochemistry to analyse sialoglycans in a TMA comparing untreated primary prostate tissue to metastatic CRPC growing in bone (TMA cohort 4). Our findings show that ligands for all three Siglecs are increased in treatment resistant tumours growing in bone relative to untreated primary prostate cancer (Figure 2A). Next, we further analysed Siglec-7 ligands in three additional TMA cohorts including unique and richly annotated clinical tissues. Analysis of a TMA constructed as part of the Movember Global Action Plan 1 Unique tissue microarray (GAP1-UTMA) project [35] (TMA cohort 5), revealed Siglec-7 ligands are expressed at similar levels in untreated (hormone naive) primary prostate tumours compared to treatment resistant (CRPC) tissues (n=161, unpaired t test, p=0.0904) (Figure 2B). Next, to assess the heterogeneity of Siglec-7 ligands across different anatomic sites in lethal prostate cancer metastasis, we analysed a 160 case TMA, also constructed as part of the Movember GAP1-UTMA project [35], that contains primary prostate tissue and rapid autopsy tissue obtained from lethal visceral and bone metastatic tumours (TMA cohort 6). Analysis of this TMA confirmed upregulation of Siglec-7 ligands in prostate cancer tissue relative to matched normal tissue from the same patient (Supplementary Figure 2) and further revealed significant upregulation of Siglec-7 engaging sialoglycans in bone metastatic prostate tumours (relative to unmatched primary prostate tissue or matched visceral tumours from the same patient) (Figure 2C). Finally, to assess if the expression of Siglec-7 ligands might correlate with survival times in prostate cancer patients, we analysed a previously published TMA comprising 100 cases of primary prostate tissues [36] (TMA cohort 7). Here, when we stratified prostate cancer patients based on high and low HYDRA-7 staining levels, defined as the top and bottom 50^th^ percentile of expression, patients with high HYDRA-7 levels had significantly poorer survival rates compared to patients with low HYDRA-7 levels (Figure 2D). In summary, these data show that the expression of sialoglycan ligands for Siglec-7 correlates with bone metastasis and poorer prognosis in patients with prostate cancer.

**Figure 2.**
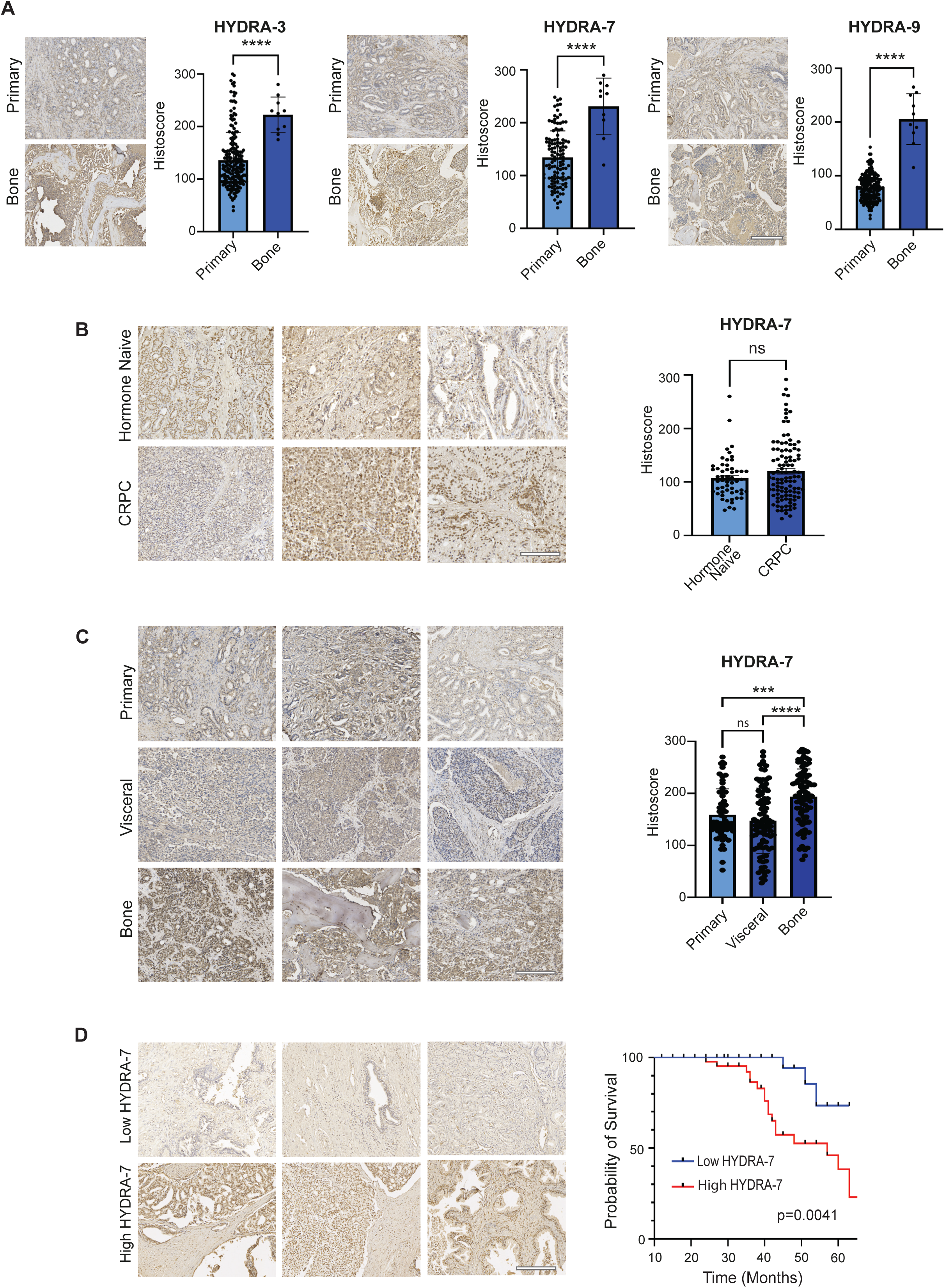
Siglec-engaging tumour immunosuppressive glyco-codes are upregulated in aggressive prostate tumours and correlate with poor prognosis. (**A**) HYDRA immunohistochemistry analysis of ligands for Siglec-3, Siglec-7, and Siglec-9 in untreated primary prostate tissue compared to metastatic castrate resistant cancer (CRPC) growing in bone suggests all three sialoglycan ligands are increased in treatment resistant metastatic tumours relative to untreated primary prostate cancer tissues (n=205, unpaired t tests, HYDRA-3 p <0.0001, HYDRA-7 p<0.0001, HYDRA-9 p<0.0001). Scale bar is 200□µm. (**B**) Analysis of Siglec-7 ligands in a TMA generated by the Movember Global Action Plan 1 Unique tissue microarray (GAP1-UTMA) project [35]. HYDRA-7 immunohistochemistry shows Siglec-7 ligands are expressed at similar levels in untreated / hormone naïve primary prostate tumours compared to therapy resistant (CRPC) tissues (n=161, unpaired t test, p=0.0904). Scale bar is 200□µm. (**C**) HYDRA immunohistochemistry analysis of Siglec-7 ligands in a TMA containing primary prostate tissue and rapid autopsy tissue obtained from lethal visceral and bone metastatic tumours [67]. HYDRA-7 Histoscores were significantly higher in lethal bone metastatic prostate tumours compared to unmatched primary prostate tumours (n=238, Welsh’s ANOVA test, p<0.0001). The levels of Siglec-7 ligands were significantly higher in prostate derived tumours growing in bone compared to matched visceral tumour tissue from the same patient (n=100, paired t test, p<0.0001). Scale bar is 200□µm. (**D**) HYDRA-7 immunohistochemistry analysis of Siglec-7 ligands in a 100-case prostate cancer TMA. Stratification of patients based on high and low Siglec-7 ligand levels shows patients with high HYDRA-7 levels (defined as the top 50^th^ percentile of expression) had significantly poorer survival rates compared to patients with low HYDRA-7 levels (defined as the bottom 50^th^ percentile of expression) (n=100, Kaplan-Meier regression model, p= 0.0041). Scale bar is 200□µm.

### 3. Siglec receptors are expressed by immune cells in the prostate cancer bone metastatic TIME

Our findings above show that immunosuppressive sialoglycans are upregulated in prostate tumours and higher levels of these glycans correlate with metastasis and poorer patient prognosis. Recent findings show that Siglec-7 and -9 are mainly expressed by myeloid cells in both primary and bone metastatic prostate tumours [27, 32]. However, no comprehensive studies have been reported that integrate both Siglec-ligand staining with direct assessment of Siglec receptors on immune cells in the same tissues. Therefore, to monitor the corresponding patterns of Siglec expression at single cell resolution in prostate cancer, we analysed previously published single cell RNA-sequencing data from: 1) primary prostate tumours [37], and 2) prostate-derived tumour metastases growing in bone [38], and further validated our findings by performing dual immunofluorescence analysis of whole slide serial sections of independent clinical tissue. Our findings confirm Siglec-7 and -9 are expressed mainly on myeloid cells in clinical prostate cancer tissue and provide further insight into the expression of other Siglec receptors in prostate tumours. We show that Siglec-7, -9, -10 and -15 are co-expressed with the myeloid marker CD14 and the M2 macrophage marker CD206 in both primary and bone metastatic tumours that also express immunosuppressive sialoglycans (Figure 3 and Supplementary Figures 3-6). Furthermore, we reveal that Siglec-2 is co-expressed with the B cell marker CD20, and in bone metastatic tissue, Siglec-3 and - 15 are expressed with the osteoclast marker cathepsin K. Siglec receptors co-localised with α-methylacyl-CoA racemase (AMACR), confirming their expression within cancerous tissues. Together with the findings above, our data provides evidence of Siglec-engaging sialoglycans and their respective immunoreceptors in prostate-derived tumours growing in bone and suggests that Siglec biology may play a role in immune cell functions in both primary and bone metastatic prostate cancer.

**Figure 3.**
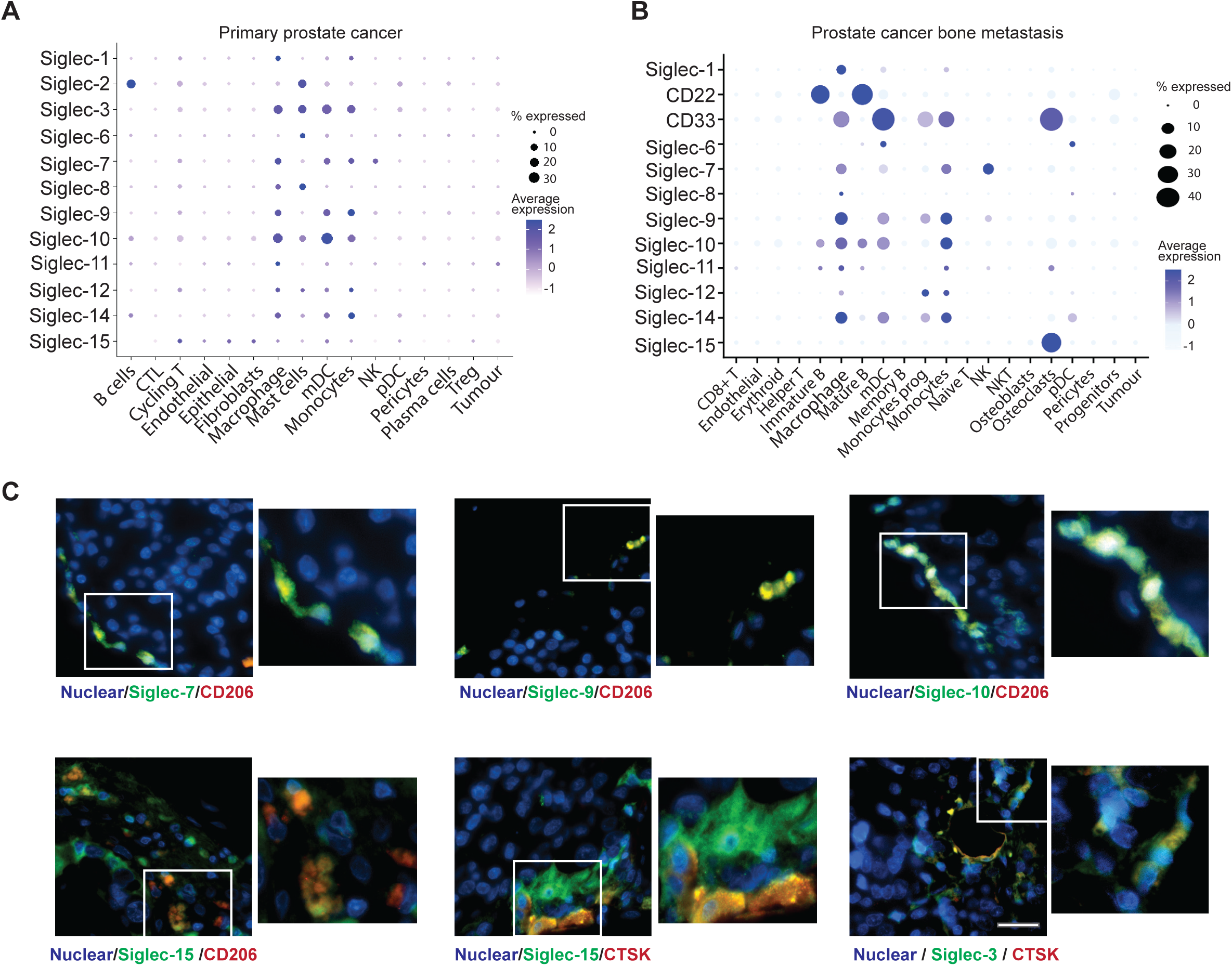
Siglec receptors are expressed by infiltrating immune cells in prostate-derived tumours growing in bone. Dot plots of *Siglec* gene expression levels at single cell resolution in prostate tumours using previously published single cell RNA-sequencing data from (**A**) primary prostate tumours [37] and (**B**) prostate cancer bone metastatic tissues [38]. (**C**) Dual immunofluorescence analysis of Siglec receptors and immune cell markers in independent prostate cancer bone metastasis tissue samples. This analysis was also carried out for primary prostate tissue (Supplementary Figure 4). Siglec-7, -9, -10 and -15 are co-expressed with CD14 (a myeloid marker) and CD206 (an M2 macrophage marker) in bone metastatic tumours. Furthermore, we reveal Siglec-3 and -15 are expressed with cathepsin K (an osteoclast marker). Scale bar is 20□µm.

### 4. An engineered bisialidase (E-612) can effectively strip Siglec ligands from prostate cancer cells

Our findings above correlate the expression of Siglecs on immune cells with Siglec-engaging sialoglycans on prostate cancer cells with bone metastasis and reduced survival times in patients. We thus hypothesised that removing sialoglycans from prostate cancer cells could be a novel therapeutic option to treat advanced prostate cancer. To investigate this hypothesis, we evaluated the potential of a proprietary engineered bisialidase reagent termed E-612, developed by Palleon Pharmaceuticals, to strip sialoglycans from prostate cancer cells. We used flow cytometry and immunofluorescence to assess the impact of E-612 treatment on prostate cancer cell recognition by several lectins. On CWR22Rv1 and murine RM1 prostate cancer cells, E-612 treatment led to reduced staining with α-2,3 Lectenz [39] that recognizes α-2,3-linked sialic acid residues (Fig. 4A, C). Concomitantly, E-612 treatment led to increased recognition by peanut agglutinin (PNA) (Fig. 4B, D) and Erythrina Cristagalli Lectin (ECL) (Supplementary Fig. 7A,B) that bind terminal galactose and lactose/LacNAc residues and therefore signify the loss of sialic acid [40]. Staining with concanavalin A (ConA) lectin revealed no difference in α-D-mannose content of E-612-treated samples and confirmed the sialic acid specificity of these changes (Supplementary Fig. 7C, D)[40]. Next, to further assess the capacity of E-612 to desialylate prostate cancer cells, we performed HYDRA-7 and HYDRA-9 flow cytometry analysis of CWR22Rv1 prostate cancer cells treated with E-612. Treatment with 1500 nM E-612 for 24 hours can remove sialic acid from the surface of CWR22Rv1 prostate cancer cells to remove ligands for both Siglec-7 and Siglec-9 (Figure 4E,F). These findings demonstrate that the bisialidase E-612 can effectively strip sialic acid from the surface of prostate cancer cells to remove immunosuppressive sialoglycans that are engaged by Siglec receptors.

**Figure 4.**
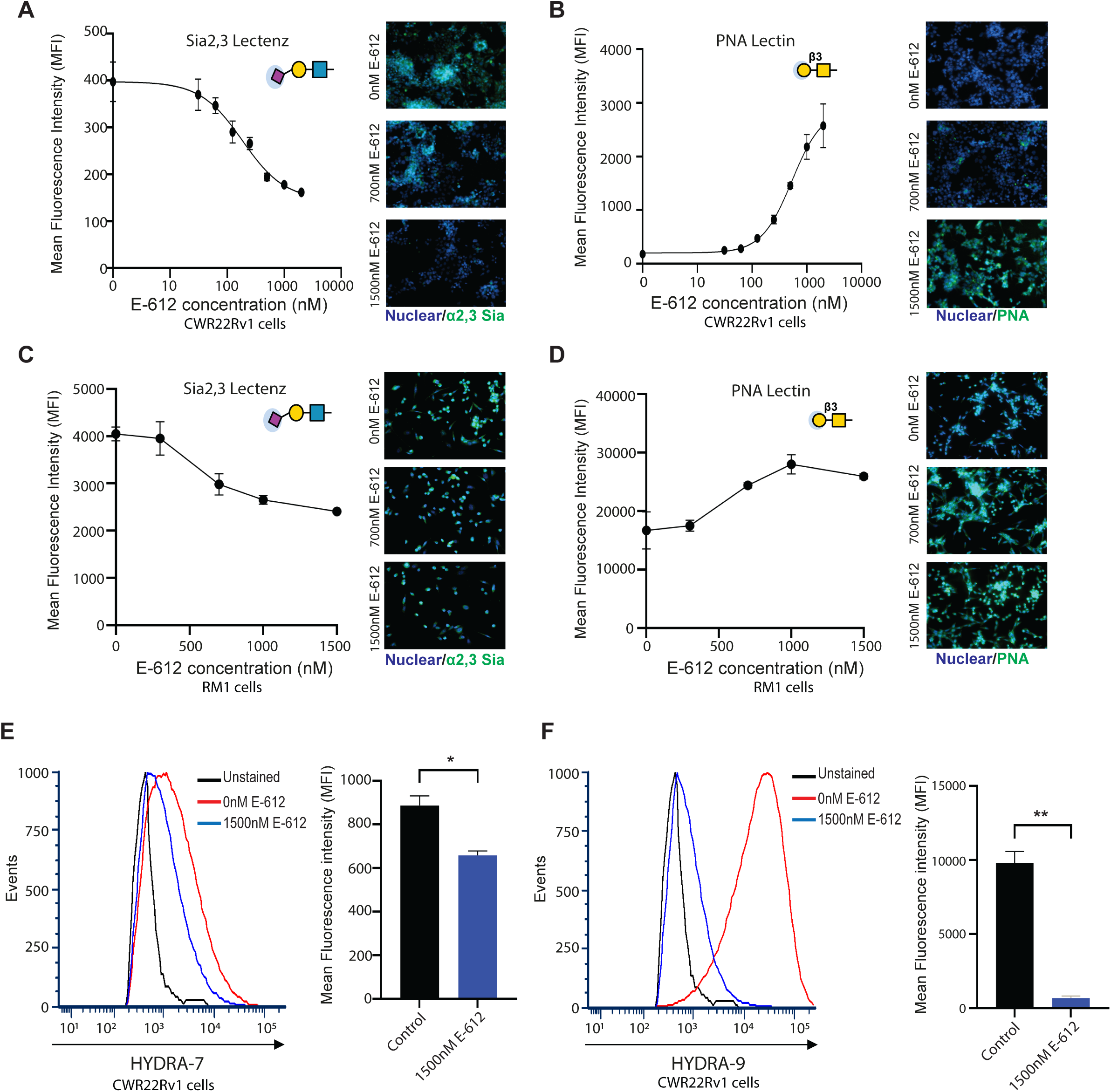
E-612 bisialidase can effectively strip Siglec ligands from prostate cancer cells. (**A**) Detection of α2-3 linked sialylated glycans in CWR22Rv1 cells using α2-3 Lectenz flow cytometry. CWR22Rv1 cells were treated with a range of concentrations of E-612 for 24 hours. CWR22Rv1 treated with 1500nM E-612 had reduced levels of α2-3 Lectenz binding indicating a reduction in α2-3 linked sialylation in these cells. Reduced levels of α2-3 linked sialylated glycans were also detected in cells treated with E-612 using α2-3 Lectenz immunofluorescence. (**B**) Lectin flow cytometry and lectin immunofluorescence assays show CWR22Rv1 cells treated with E-612 for 24 hours have increased levels of binding to PNA lectin (which recognises galactosyl residues [40] uncovered by the removal of sialic acids). (**C**) Detection of α2-3 linked sialylated glycans in mouse RM1 prostate cancer cells using α2-3 Lectenz flow cytometry. RM1 cells were treated with a range of concentrations of E-612 for 24 hours. RM1 cells treated with 1500nM E-612 had reduced levels of α2-3 Lectenz binding indicating a reduction in α2-3 linked sialylation in these cells. Reduced levels of α2-3 linked sialylated glycans were also detected in cells treated with E-612 using α2-3 Lectenz immunofluorescence. (**D**) Lectin flow cytometry and lectin immunofluorescence assays show RM1 cells treated with E-612 for 24 hours have increased levels of binding to PNA lectin (which recognises galactosyl residues [40] uncovered by the removal of sialic acids). (**E**) HYDRA-7 flow cytometry shows treatment with 1500nM E-612 significantly reduces the binding of HYDRA-7 reagent to CWR22Rv1 cells indicating that E-612 has removed the sialoglycan ligands for Siglec-7 from the cell surface (unpaired t test, p=0.0207) (**F**) HYDRA-9 flow cytometry shows treatment with 1500nM E-612 significantly reduces the binding of HYDRA-9 reagent to CWR22Rv1 cells indicating that E-612 has removed the sialoglycan ligands for Siglec-9 from the cell surface (unpaired t test, p=0.0058).

### 5. Therapeutic desialylation with E-612 reduces the growth of tumours and prolongs survival times of mice with prostate cancer bone metastasis

Previous studies have shown that targeting the ligands for Siglec-7/9 by knockout of the glycosyltransferase ST3GAL1 or by using Siglec-7/9 blocking antibodies can decrease prostate tumour growth and enhance anti-tumour immunity [27, 32]. We have previously shown that pre-treatment of prostate cancer cells with a sialyltransferase inhibitor leads to a reduction of sialylated glycans and suppresses bone metastasis [26]. These findings raise the possibility that systemic therapies to disrupt the Siglec-sialoglycan axis will likely be effective against bone metastatic prostate cancer. However, this has not yet been tested using pre-clinical mouse models. As Siglec-E is the most broadly expressed inhibitory Siglec in mice [41] and has previously been identified as a mediator of the effects of therapeutic desialylation [42], we next tested if Siglec-E is expressed in the mouse bone metastatic TIME. Analysis of PC3 prostate tumours growing within the tibias of mice revealed that Siglec-E is expressed by myeloid cells within the bone TIME (Figure 5A). This finding raised the possibility of targeting the sialoglycan-Siglec-E axis to treat bone metastatic prostate cancer in mice. Next, we utilised syngeneic mouse models to investigate whether desialylation using E-612 can be used therapeutically for metastatic prostate cancer.

**Figure 5.**
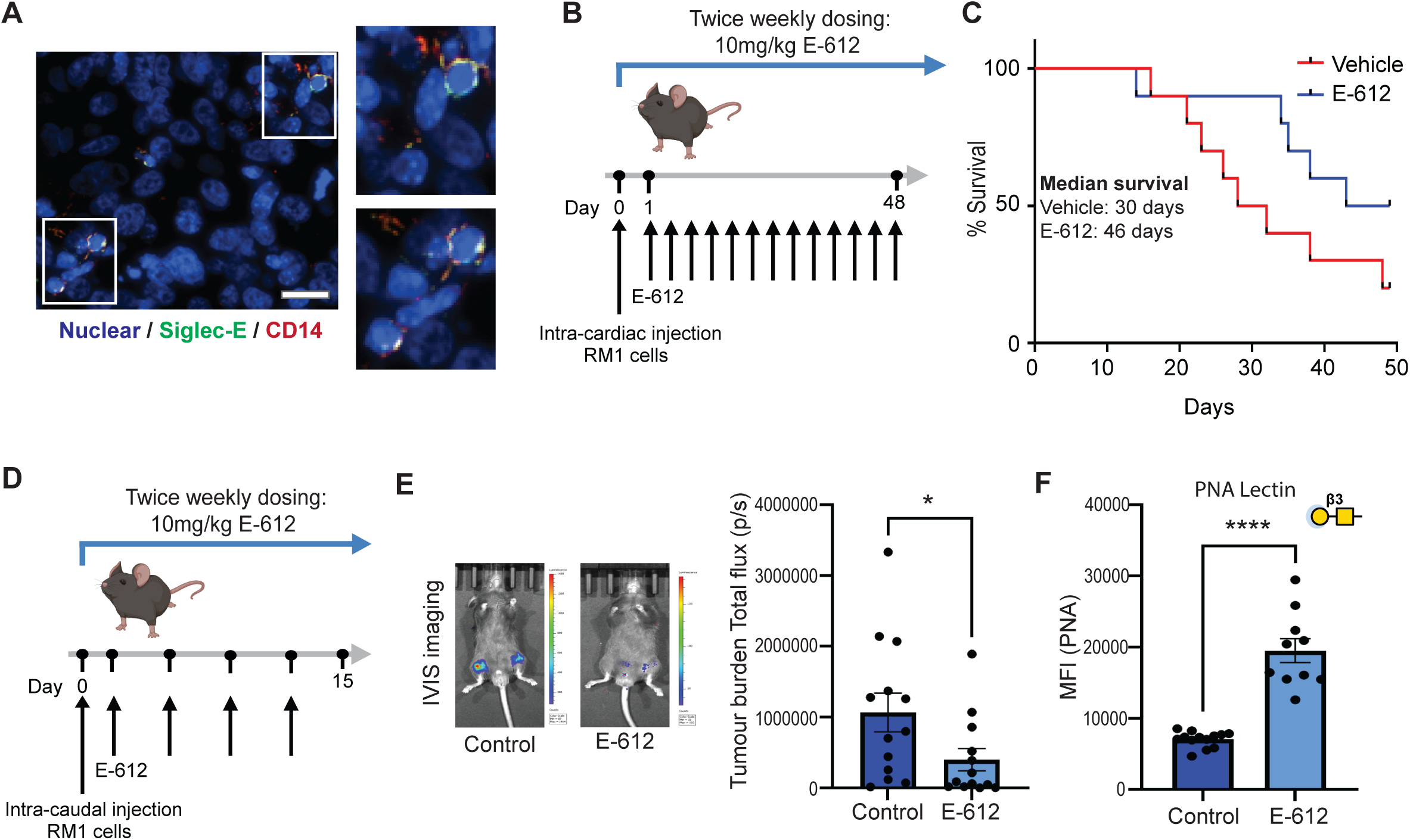
Desialylation of prostate cancer cells with E-612 reduces the growth of tumours and prolongs survival times of mice with metastasis. (**A**) Dual immunofluorescence analysis of Siglec-E and CD14 (a myeloid marker) in PC3 prostate cancer tumours growing in mouse tibias. Siglec-E is co-localised with CD14 confirming its expression by myeloid cells within the murine prostate cancer TIME. Scale bar is 20□µm. (**B**) Using an RM1 intra-cardiac injection metastasis model, we investigated if therapeutic desialylation by E-612 can increase survival times of mice with metastatic prostate cancer. (**C**) Twice weekly dosing with 10mg/kg E-612 via intra-peritoneal injection prolongs the median survival rates of mice with metastatic prostate cancer by 53%. (**D**) Using an RM1 intra-caudal injection bone metastasis model [43] we tested if E-612 mediated tumour desialylation can suppress the growth of bone metastatic prostate tumours. (**E**) Systemic treatment with E-612 (twice weekly dosing with 10mg/kg E-612 via intra-peritoneal injection) significantly reduces prostate cancer bone metastasis tumour burden (unpaired t test, p=0.0456). (**F**) PNA lectin flow cytometry analysis of white blood cells from mice treated with E-612. Systemic E-612 therapy significantly increases the levels PNA lectin binding to circulating immune cells (PNA lectin recognises galactosyl residues that are uncovered by the removal of sialic acids [40]) (Welch’s t test, p<0.0001).

Consistent with this, using an RM1 intra-cardiac injection metastasis model, systemic treatment with E-612 dosed with 10 mg/kg twice per week via intra-peritoneal injection prolonged the median survival rates of mice with metastatic prostate cancer by 53% (Figure 5B,C). Next, using an RM1 intra-caudal injection bone metastasis model, that recapitulates the process of bone metastasis [43], we tested if E-612 mediated tumour desialylation can suppress the growth of bone metastatic prostate tumours. Systemic treatment with E-612 significantly reduced the tumour burden of bone metastatic prostate cancer (unpaired t test, p=0.0456) (Figure 5D, E). In both these studies, the body weights of both treatment groups increased similarly over the course of the study suggesting no signs of acute toxicity. To confirm that E-612 can effectively mediate desialylation *in vivo*, we used PNA lectin flow cytometry to monitor the levels of sialic acid on white blood cells isolated from mice undergoing E-612 therapy. Our data revealed that systemic E-612 treatment effectively removes sialic acid from circulating immune cells (Figure 5F). These findings demonstrate the feasibility of using therapeutic *in vivo* desialylation to suppress the growth of bone metastatic prostate cancer.

## DISCUSSION

Prostate cancer is a leading cause of cancer-related mortality in men over the age of 50 [44]. Although effective therapies for prostate cancer exist, resistance to therapy is very common, and there is a continuing need to develop new treatment options, especially for prostate cancer that has spread to bone. Many studies have profiled the molecular changes correlating with prostate cancer progression and bone metastasis, with most research focused on changes to the genome, transcriptome, epigenome, proteome and metabolome. However, despite the functional link between aberrant glycosylation and all of the cancer hallmarks [12], and the prostate being an abundant secretor of glycoproteins [45], the glycome is relatively understudied in prostate cancer [46]. Novel therapies targeting sialoglycans are currently being developed and trialled, and it is timely to investigate how prostate cancer patients could best benefit from these advances.

Here, we utilise three novel reagents (HYDRA-3, -7, and -9) [33, 47] to investigate sialoglycan ligand expression across the full clinical heterogeneity of clinical prostate tumour tissue. We reveal that sialoglycans that engage Siglec-3, -7 and -9 are upregulated in prostate tumours. Furthermore, we show sialoglycan ligands for Siglec-7 correlate with bone metastasis and reduced survival times in prostate cancer patients. Analysis of clinical tissue shows Siglec receptors (which engage immunosuppressive sialoglycans) are expressed by immune cells in the bone metastatic prostate tumour immune microenvironment (TIME), providing evidence that this axis plays a role in immune cell functions in bone metastatic prostate cancer. We demonstrate that an engineered dual-action sialidase (E-612) can effectively remove sialic acid from the surface of prostate cancer cells, and that systemic treatment with E-612 impedes the growth of prostate tumours in bone and improves survival times in mice. These findings identify a novel mechanism (the Siglec-sialoglycan axis) as important in the growth of prostate-derived tumours in bone, demonstrate that this axis is clinically actionable, and provide a rationale for further investigation of this biology to develop new therapies for advanced prostate cancer.

Aberrant sialylation has been understudied in prostate cancer, primarily due to the lack of useful reagents for detecting this biology in tissue [21]. Our study utilises novel reagents to comprehensively monitor Siglec-engaging immunosuppressive sialoglycans across the full heterogeneity of clinical prostate cancer tissue. Our findings correlate the expression of Siglec-engaging sialoglycans on prostate cancer cells with bone metastasis, thus suggesting that Siglec biology may play a role in immune cell functions in bone metastatic prostate cancer and revealing the potential importance of the Siglec-sialoglycan axis in the bone TIME. Macrophages are the most abundant immune cell types in prostate cancer and studies show CD206+ macrophages are linked to poor patient prognosis and are uniquely enriched in the bone tumour microenvironment of prostate cancer [38, 48–50]. Specifically, we show that in bone metastatic tumours Siglec-7, -9, -10 and -15 are expressed by M2 macrophages, and Siglec-15 is expressed by osteoclasts. Therapies aimed at reducing M2 macrophage infiltration, or promoting macrophage repolarisation to pro-inflammatory M1 phenotypes, are being actively investigated for metastatic CRPC [51], and it will be important for future studies in this area to consider Siglec-sialoglycan biology. Siglec-15 has a well-established role in osteoclast differentiation [52–54], and in the context of bone metastasis, Siglec-15 can contribute to the ‘vicious cycle’ of bone destruction and tumour growth [55].

The Siglec-15 sialoglycan axis was recently identified as a novel immune checkpoint in breast cancer that can be targeted using therapeutic antibodies to inhibit bone metastasis [55], thus raising the possibility that this newly discovered immunological pathway could likely also be targeted for prostate cancer therapy.

Our study identifies therapeutic desialylation using E-612 as a promising therapeutic strategy against prostate tumours, and specifically for prostate-derived tumours growing in bone, which represent an incurable major clinical burden. While initial interrogation of the Siglec-sialoglycan axis in cancer focused on blocking Siglec receptors [16], it has been suggested that, due to redundancy, targeting a single Siglec or one or more of its ligands, may not be an effective means of drugging this axis. Engineered enzymes to strip terminal sialic acids from the glycans that cover the surface of cancer and immune cells could overcome Siglec redundancy to effectively inhibit multiple Siglec receptors. Bisialidase enzymes have been used therapeutically in mouse models to halt the growth of breast, melanoma, and colon cancers [56]. Our findings indicate that this therapeutic approach is also therapeutically relevant for prostate cancer, and specifically for patients with bone metastasis. Targeted sialidase therapy, with a ligand that binds to HER2 receptors has been developed and has demonstrated efficacy to treat breast cancer [57, 58]. We propose that a similar therapeutic strategy could be applied to prostate cancer by using prostate-specific membrane antigen (PSMA) or prostate stem cell antigen (PSCA) to leverage enzymatic sialoglycan degradation to specifically target prostate cancer cells [32].

Numerous studies have highlighted a role for aberrant sialylation in cancer therapy resistance, suggesting that targeting cancer-associated glycans may enhance responsiveness to cancer treatment [16, 42, 59, 60]. A recent breakthrough study demonstrated that therapeutic desialylation can reshape macrophage phenotypes to allow effective immune checkpoint blockade (ICB) [42]. ICB for the treatment of advanced prostate cancer is effective in only a minority of patients [61]. Ongoing clinical trials are now focused on developing novel combination therapies aimed at sensitising prostate tumours to immune checkpoint blockade [62–65], and moving forward, once additional pre-clinical evaluation studies have been performed, therapeutic desialylation in combination with ICB offers a promising novel therapeutic avenue for prostate cancer to explore. Our recent findings suggest sialylation inhibition can partially revert acquired resistance to enzalutamide in prostate cancer cells [60], providing rationale to further investigate therapeutic desialylation in combination with hormone therapy as a novel therapeutic approach in patients with advanced prostate cancer.

In conclusion, our study reveals Siglec-engaging immunosuppressive sialoglycans are upregulated in prostate tumours and correlate with bone metastasis and reduced survival times in patients. We propose that sialoglycan-Siglec biology is important in the tumour microenvironment of bone metastatic prostate cancer and demonstrate that therapeutic targeting of tumour sialylation can effectively impede the growth of prostate cancer in bone. Together, these results provide strong rationale for the further clinical development of sialoglycan-Siglec targeting therapies for bone metastatic prostate cancer, and their investigation in combination with existing immunotherapy and anti-androgen therapies.

## METHODS

### Clinical cohorts

**TMA cohort 1**: 40 case human prostate cancer TMA was purchased from Novus Bio (NBP2-30169).

**TMA cohort 2**: A previously published 96 case TMA comprising cores of normal prostate tissue and prostate adenocarcinoma of different Gleason grades was purchased from US Biomax (PR1921b) [66].

**TMA cohort 3**: A previously published TMA consisting of prostatectomy tissue from 200 cases of prostate cancer and matched normal tissue (four cores each) [25, 66].

**TMA cohort 4**: A previously published TMA consisting of prostatectomy tissue from 200 cases of untreated primary prostate cancer and 10 cases of rapid autopsy FFPE tissue samples from prostate-derived tumours growing in bone [26].

**TMA cohort 5**: 174 case TMA with untreated (hormone naïve) and treatment resistant (CRPC) tissues constructed as part of the Movember Global Action Plan 1 Unique tissue microarray (GAP1-UTMA) project [35].

**TMA cohort 6**: 100 case CRPC metastatic clinical heterogeneity TMA intended to assess the heterogeneity of molecular markers across different anatomic sites in lethal prostate cancer metastasis. This TMA has been previously published TMA [67] was created as part of the Movember Global Action Plan 1 Unique tissue microarray (GAP1-UTMA) project [35].

**TMA cohort 7**: 100 case prostate cancer and normal tissue TMA with survival data was purchased from US Biomax (MPR1005sur) [36].

### Primary prostate cancer tissue

Patient tissue samples were collected with ethical permission from Castle Hill Hospital (Cottingham, Hull) (Ethics Number: 07/H1304/121). Use of patient tissue was approved by the Local Research Ethics Committees. Patients gave informed consent and all patient samples were anonymized.

### Bone metastasis tissue samples from rapid autopsy

20 cases of rapid autopsy FFPE tissue samples from prostate-derived tumours growing in bone were kindly provided by Dr Colm Morrissey (University of Washington) via the Prostate Cancer Biorepository Network (PCBN). Biopsies of metastatic bone sites were obtained from patients with CRPC within hours of death using a cordless drilling trephine (DeWalt Industrial Tool) and model 2422-51-000 trephine (DePuy). Bone cores were fixed in 10% neutral buffered formalin, decalcified with 10% formic acid and paraffin embedded.

### Immunohistochemistry

The immunohistochemistry (IHC) protocol using the HYDRA reagents adapted from previously published methodology [22]. HYDRA-3, HYDRA-7 and HYDRA-9 reagents, which correspond to ligand binding domains of their corresponding Siglecs (Siglec-3, Siglec-7 and Siglec-9 respectively) were provided by Palleon Pharmaceuticals (Waltham, MA). HYDRA reagents were diluted in Antibody Diluent (Dako, S302283-2). All slides were stained using the IHC Prep & Detect Kit for Rabbit/Mouse Primary Antibody (Proteintech, PK10019), following the manufacturer’s protocols. A Sodium Citrate Antigen Retrieval Buffer (50X) (Proteintech, PR30001) was used, and tissues were incubated with the HYDRA-3 at 4.5 μg/mL, HYDRA-7 at 1.5 μg/mL or HYDRA-9 at 0.25 μg/mL for 1 hour before the secondary antibody incubation. HistoClear (National Diagnostics, HS-200) and HistoMount (National Diagnostics, HS-103) were used instead of Xylene and the mounting media from the kit. The TMAs were scored by a pathologist using the 0–300 HistoScore method [68, 69]. Only epithelial cells were scored. Images were acquired and processed using the ZEISS Axioscan 7 and ZEN Microscopy Software (blue edition).

### Analysis of Single cell RNAseq data

Primary prostate cancer and prostate cancer bone metastases single cell data were downloaded from GSE143791 [38] and GSE181294 [37]. Normalized gene expression matrixes and the cell annotations were utilised. Visualisation of the Single-cell RNAseq data was performed using the Seurat package (v 5.3.0) in RStudio. Standard Seurat workflows were used for visualization of selected marker gene expression using the DotPlot function.

### Cell culture

CWR22RV1 (RRID:CVCL_1045) and RM1 cells (RRID:CVCL_B459) were cultured as described previously [26, 70]. All of the cell lines used were validated using STR profiling and tested monthly for mycoplasma contamination.

### Immunofluorescence (clinical tissue and cell lines)

To monitor sialylation changes following E-612 treatment, cells were cultured in Nunc™ Cell-Culture Treated 4-well dishes (Thermo Scientific, 176740) in complete media and treated with E-612 (Palleon Pharmaceuticals, Waltham, MA) for 24 hrs at 37 °C in a humidified incubator with 5% carbon dioxide. Next, cells were washed with PBS before permeabilization and fixation with ice-cold absolute methanol for 10 min at −20 °C. After another PBS wash, cells were blocked with 1X Carbo-Free™ Blocking Solution (1X CFB) (Vector Laboratories, SP-5040-125) for 1 hr at room temperature, before they were incubated overnight in the dark at 4 °C in 1:1000 Fluorescein-labelled Peanut Agglutinin (PNA) Lectin (Vector Laboratories, FL-1071), 1:1000 Fluorescein-labelled Erythrina Cristagalli Lectin (ECL, ECA) (Vector Laboratories, FL-1141-5), 1:1000 Fluorescein-labelled Concanavalin A (Con A) Lectin (Vector Laboratories, FL-1001-25) or 1:400 SureLight® 488-conjugated SiaFind Alpha 2,3-Specific Lectenz® (Lectenz Bio, SK2301F). Finally, following PBS washes, cells were stained with 1:1200 Hoechst Nucleic Acid Stain (Thermo Scientific, 62249) for 15 min at room temperature. Cells were then washed with PBS and mounted using ProLong™ Gold Antifade Mountant (Thermo Fisher, P36930).

For Siglec dual staining, FFPE-fixed prostate clinical tissues were dewaxed and rehydrated before antigen retrieval at 95 °C for 20 min using a Sodium Citrate Antigen Retrieval Buffer (50X) (Proteintech, PR30001). Tissues were then blocked with 1X CFB for 1 hr at room temperature, following overnight incubation in the dark at 4 °C in 1:400 CD22 Monoclonal Antibody (Proteintech, 66103-1-Ig), 1:200 Anti-CD33 Antibody [WM53] (Abcam, ab252263), 1:200 Siglec-7/CD328 Polyclonal Antibody (Proteintech, 13939-1-AP), 1:200 Siglec-9 Polyclonal Antibody (Proteintech, 13377-1-AP), 1:100 Siglec-10 Polyclonal Antibody (Invitrogen, PA5-55501), 1:100 Siglec-15 Polyclonal Antibody (Abcam, ab198684), 1:50 Mouse Siglec-E Antibody (R&D Systems, AF5806), 1:200 CD14 Rabbit Polyclonal Antibody (Proteintech, 17000-1-AP), 1:200 CD14 Mouse Monoclonal Antibody (Proteintech, 60253-1-Ig), 1:500 AMACR/p504S Rabbit Polyclonal Antibody (Proteintech, 15918-1-AP), 1:200 AMACR Mouse Monoclonal Antibody [UMAB68] (UltraMAB, UM870012), 1:200 CD206 Rabbit Polyclonal Antibody (Proteintech, 18704-1-AP), 1:200 CD206 Mouse Monoclonal Antibody (Proteintech, 60143-1-Ig), 1:200 Cathepsin K Rabbit Polyclonal Antibody (Proteintech, 11239-1-AP), 1:200 Cathepsin K Mouse Monoclonal Antibody [E-7] (Santa Cruz, sc-48353), 1:200 CD20 Rabbit Polyclonal Antibody (Proteintech, 24828-1-AP) or 1:100 Pan Cytokeratin Monoclonal Antibody (AE1/AE3), Alexa Fluor™ 488-conjugated (Invitrogen 53-9003-80). Next, tissues were washed with PBS before 1 hr incubation in the dark at room temperature in 1:1000 Goat Anti-Rabbit IgG H&L (Alexa Fluor® 594) (Abcam, ab150080), 1:1000 Goat Anti-Mouse IgG H&L (Alexa Fluor® 488) (Abcam, ab150113), or 1:500 Multi-rAb® CoraLite® Plus 594-Goat Anti-Rabbit Recombinant Secondary Antibody (H+L) (Proteintech, RGAR004) and 1:100 Fluorescein (FITC)–conjugated Donkey Anti-Goat IgG(H+L) (Proteintech, SA00003-3). Finally, following PBS washes, tissues were stained with 1:1200 Hoechst Nucleic Acid Stain (Thermo Scientific, 62249) for 15 min at room temperature. Tissues were then washed with PBS and mounted using ProLong™ Gold Antifade Mountant (Thermo Fisher, P36930). Images were acquired and processed with the ZEISS Axio Imager 4 and ZEN Microscopy Software (blue edition).

### Expression and purification of engineered human Neu2 sialidase-Fc fusion (E-612)

A mutant of human Neu2 sialidase, incorporating structure-guided stabilizing mutations, fused to human IgG1 Fc fragment, was designated E-612. DNA encoded E-612 was transfected into a CHO-K1 stable cell pool for recombinant protein expression (WuXi Biologics, Shanghai, China). Protein expression was conducted at 3 litre scale and cultured for 15 days following standard cell culture processes. E-612 was purified from cell culture supernatants using a 3-step process: Protein-A chromatography, followed by CHT (ceramic hydroxyapatite type II) chromatography, and lastly SEC (size exclusion chromatography).

CHT chromatography used the following process parameters, binding in CHT-A buffer (30mM Phosphate at pH6.5, 10mM MES PH6.5, 0.1 mM CaCl_2_), and gradient elution with 35% CHT-B buffer (30mM Phosphate at pH6.5, 100mM MES pH6.5, 0.1 mM CaCl_2_, 1M NaCl). Final protein was buffer exchanged into the formulation buffer (137 mM NaCl, 2.7 mM KCl, 10 mM Na_2_HPO_4_, 1.8 mM KH_2_PO_4_, pH 7.4, 25 mM Na Citrate, 1 mM CaCl_2_, 10% (w/v) Glycerol).

### Flow cytometry

Cells were cultured in 96-well plates (Sarstedt, 83.3924) in complete media and treated with E-612 (Palleon Pharmaceuticals, Waltham, MA) for 24 hrs at 37 °C in a humidified incubator with 5% carbon dioxide. Cells were then washed with PBS, trypsinised and centrifuged at 500×g for 5 min at room temperature. They were washed twice with 1X CFB before being resuspended in 100 μL of 1:2000 Fluorescein-labelled Peanut Agglutinin (PNA) Lectin (Vector Laboratories, FL-1071), 1:2000 Fluorescein-labelled Erythrina Cristagalli Lectin (ECL, ECA) (Vector Laboratories, FL-1141-5) or 1:400 SureLight® 488-conjugated SiaFind Alpha 2,3-Specific Lectenz® (Lectenz Bio, SK2301F) in 1X CFB and incubated for 30 min in the dark at 4 °C. Next, cells were washed with PBS twice before being resuspended in 500 μL of PBS with 1 μg/mL propidium iodide. Finally, cells were run on a BD LSRFortessa™ Cell Analyzer (BD Biosciences) with 10,000 events recorded per sample. Data was analysed using the FCS Express™ Flow Cytometry Analysis Software.

### Murine white blood cell fixation and flow cytometry

200 μl blood was collected from the RM1 intra-caudal study mice from terminal cardiac bleeding using a 25G needle and 1 ml syringe and added to 30 μl of 0.5 μM EDTA in 2 ml collection tubes. Red blood cells were lysed using 1X Red Blood Cell (RBC) Lysis Buffer (Invitrogen, 00-4333-57). Pellets were washed twice with 1 ml 1X PBS then fixed with 1% PFA on ice for 10 min. After two washes, cells were resuspended in 1X PBS with 1% v/v BSA and stored at 4°C. For flow cytometry, pellets were stained with 100 μl 2.5 μg/ml Fluorescein labelled Peanut Agglutinin (PNA) lectin (Vector Laboratories, FL-1071-10) diluted in 1X CFB for 30 min at 4°C in the dark. After two washes with FACS buffer (1X PBS with 1% v/v FBS), cells were stained with 100 μl 0.25 μg/ml CD45-PE antibody (BioLegend, 103105) diluted in FACS buffer for 15 min at 4°C in the dark. Cells were washed twice then resuspended in 500 μl FACS buffer. Data was acquired on a BD LSRFortessa™ Cell Analyzer (BD Biosciences) with 10,000 events recorded per sample. Cells were identified by size and granularity (FSC-A vs. SSC-A) followed by gating on CD45 (561 586/15-A vs SSC-A) to isolate leukocytes. Finally, a histogram was used to determine the mean fluorescence intensity (MFI) for PNA (488 530-30-A vs Count). Data was analysed using FCS Express™ 7 Plus version no. 7.14.0020.

### Mouse models

#### RM1 metastasis study

6-week old male C57BL/6 mice (RRID:MGI:2159769) were purchased from Charles River (Kent, UK) and housed in a controlled environment in Optimice cages (Animal Care Systems, Colorado, USA) randomly located in the cage rack with a 12 h light/dark cycle at 22°C with ad libitum water and 2018 Teklad Global 18% protein rodent diet containing 1.01% Calcium (Harlan Laboratories, UK). RM1 cells [71] were resuspended in PBS to 1 x 10^6^ cells/mL. Twenty mice received single-cell suspensions of 50,000 RM1cells/100 μL PBS via intra-cardiac injection. Mice were randomised into two groups to receive intra-peritoneal injection (IP) of E-612 (10mg/kg) or vehicle control twice weekly. Upon reaching humane endpoint, mice were euthanized to generate a survival curve over a 48-day period.

### RM1 intra-caudal study

To more focus on the bone metastasis, we adopted the intra-caudal artery injection model that recapitulates the process of bone metastasis in the hindlimbs only [43]. Briefly, sixteen 6-week-old male C57BL/6 mice were inoculated with a single-cell suspension of 1 x 10^6^ RM1 cells/100 μL PBS via intra-caudal artery injection. Mice were randomised into two groups to receive IP injection of E-612 (10mg/kg) or vehicle control twice weekly from day 1 post-tumour inoculation. Tumour progression was monitored weekly based on bioluminescence using the IVIS system for 2 weeks and the images were blinded for data analysis. Mice were euthanized on day 15 post-tumour inoculation, through exsanguination under general anaesthesia, followed by cervical dislocation.

All procedures complied with the UK Animals (Scientific Procedures) Act 1986 and were reviewed and approved by the local Research Ethics Committees of the University of Sheffield under Home Office project licence PP3267943 (Sheffield, UK).

## ACKNOWLEDGEMENTS

This work was funded by Prostate Cancer UK and the Bob Willis Fund through Research Innovation Awards [RIA16-ST2-011 and RIA21-ST2-006], the Medical Research Council [MR/R015902/1], Prostate Cancer Research and the Mark Foundation for Cancer Research (grant references 6961 and 6974). This work was supported by the Francis Crick Institute which receives its core funding from Cancer Research UK (CC2127), the UK Medical Research Council (CC2127) and Wellcome Trust (CC2127). This work is also supported by the Department of Defence Prostate Cancer Research Program, DOD Award No W81XWH-18-2-0013, W81XWH-18-2-0015, W81XWH-18-2-0016, W81XWH-18-2-0017, W81XWH-18-2-0018 and W81XWH-18-2-0019 PCRP Prostate Cancer Biorepository Network (PCBN).

---

**Supplementary Figure 1.** Sialoglycans that engage Siglec-7 and -9 are increased in high Gleason grade prostate cancer. HYDRA immunohistochemistry analysis of the 96 case TMA shown in Figure 1B showed higher expression of Siglec-7 ligands (p<0.0001) and Siglec-9 ligands (p=0.0437) in prostate tumours (Gleason grade 4-5) compared to lower grade prostate tumours (Gleason grade 1-3).

**Supplementary Figure 2.** HYDRA immunohistochemistry analysis of Siglec-7 ligands in TMA cohort 6 [67]. Siglec-7 engaging sialoglycans are significantly upregulated in primary prostate tissue compared to matched normal tissue (n=58, paired t test, p<0.0001).

**Supplementary Figure 3.** UMAP map of previously published single cell RNA-sequencing data from primary prostate tumours [37] showing individual *Siglec* expression levels across different cell types.

**Supplementary Figure 4. Siglec receptors are expressed by infiltrating immune cells in primary prostate cancer tumours.** Dual immunofluorescence analysis of Siglec receptors and immune cell markers in independent primary prostate tumour tissue samples. Siglec-2, -3, -7, -9, -10 and -15 are co-expressed with CD14 (a myeloid marker) in primary prostate tumours. Siglec receptors co-localised with α-methylacyl-CoA racemase (AMACR) confirming their expression within cancerous tissues. Scale bar is 20□µm.

**Supplementary Figure 5.** UMAP map of previously published single cell RNA-sequencing data from prostate cancer bone metastatic tissues [38]. showing individual *Siglec* expression levels across different cell types.

**Supplementary Figure 6. Siglec receptors are expressed by infiltrating immune cells in prostate derived tumours growing in bone.** Dual immunofluorescence analysis of Siglec receptors and immune cell markers in independent prostate cancer bone metastasis tissue samples. Siglec-3, -7, -9, -10 and -15 are co-expressed with CD14 (a myeloid marker). Siglec-2 is co-expressed with CD20 (a B cell marker). Siglec receptors co-localised with α-methylacyl-CoA racemase (AMACR) confirming their expression within cancerous tissues. Scale bar is 20□µm.

**Supplementary Figure 7. E-612 bisialidase can effectively strip Siglec ligands from prostate cancer cells.** (**A,B**) Lectin flow cytometry and lectin immunofluorescence assays show CWR22Rv1 and RM1 cells treated with E-612 for 24 hours have increased levels of binding to Erythrina Cristagalli Lectin (ECL) (which binds terminal galactose or LacNAc epitopes [40] uncovered by the removal of sialic acids). (**C**) Staining with ConA lectin (which recognises glycans containing α-D-mannose [40] confirmed the sialic acid specificity of these changes.

