## Supplementary Figure 1-7 for "Siglec-engaging immunosuppressive sialoglycans are upregulated in prostate cancer and are targetable to suppress bone metastasis"

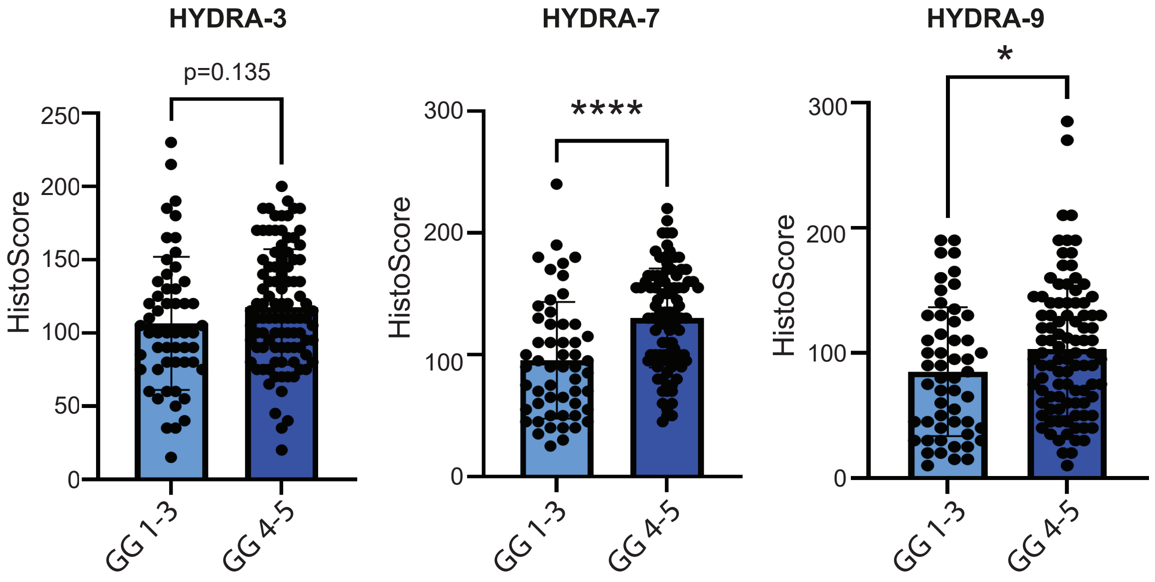

Supplementary Figure 2

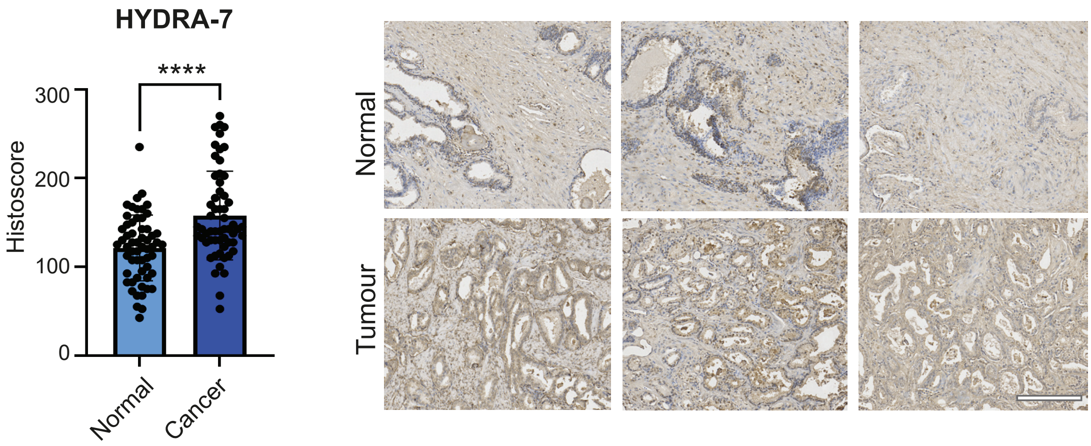

Supplementary Figure 3

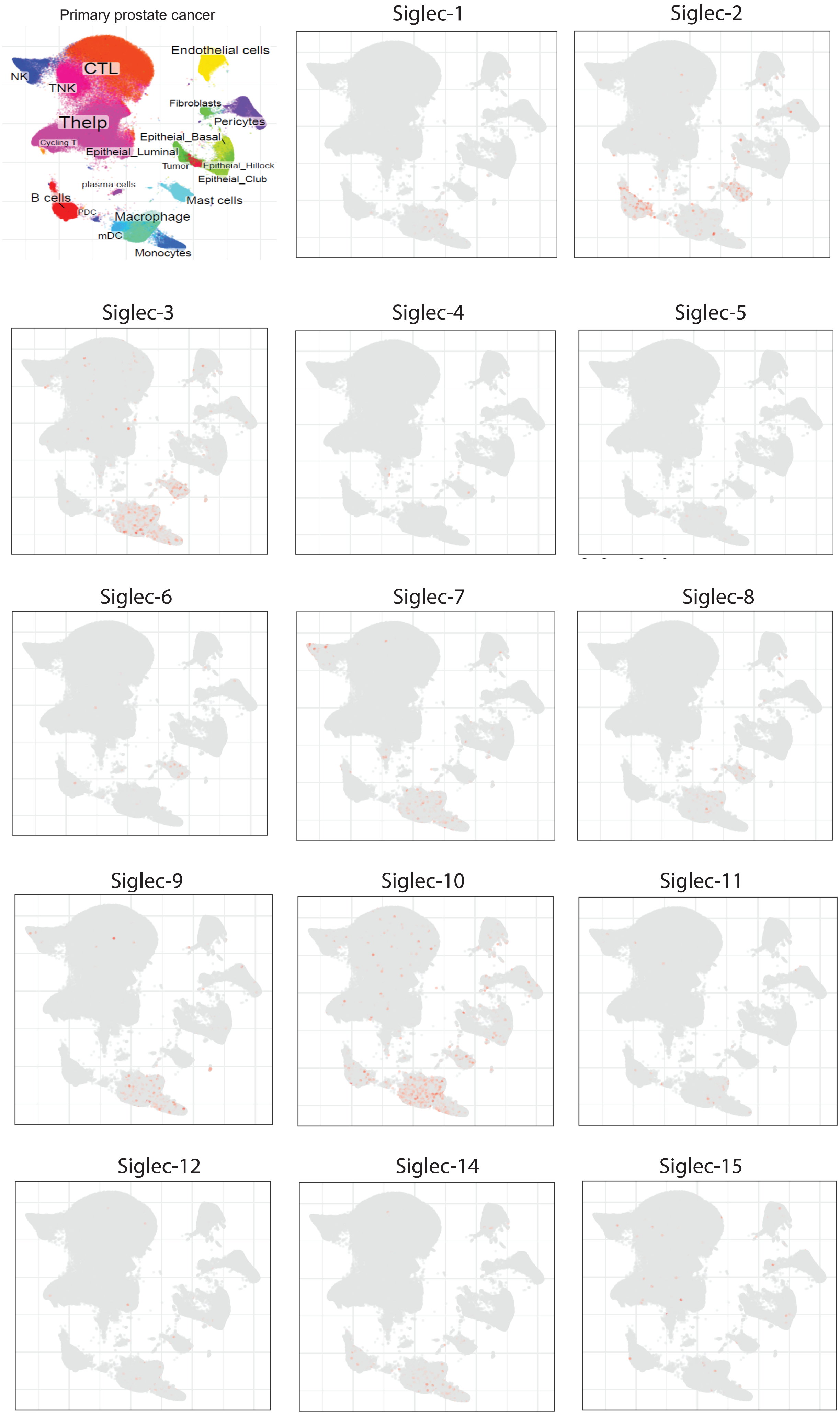

Supplementary Figure 4

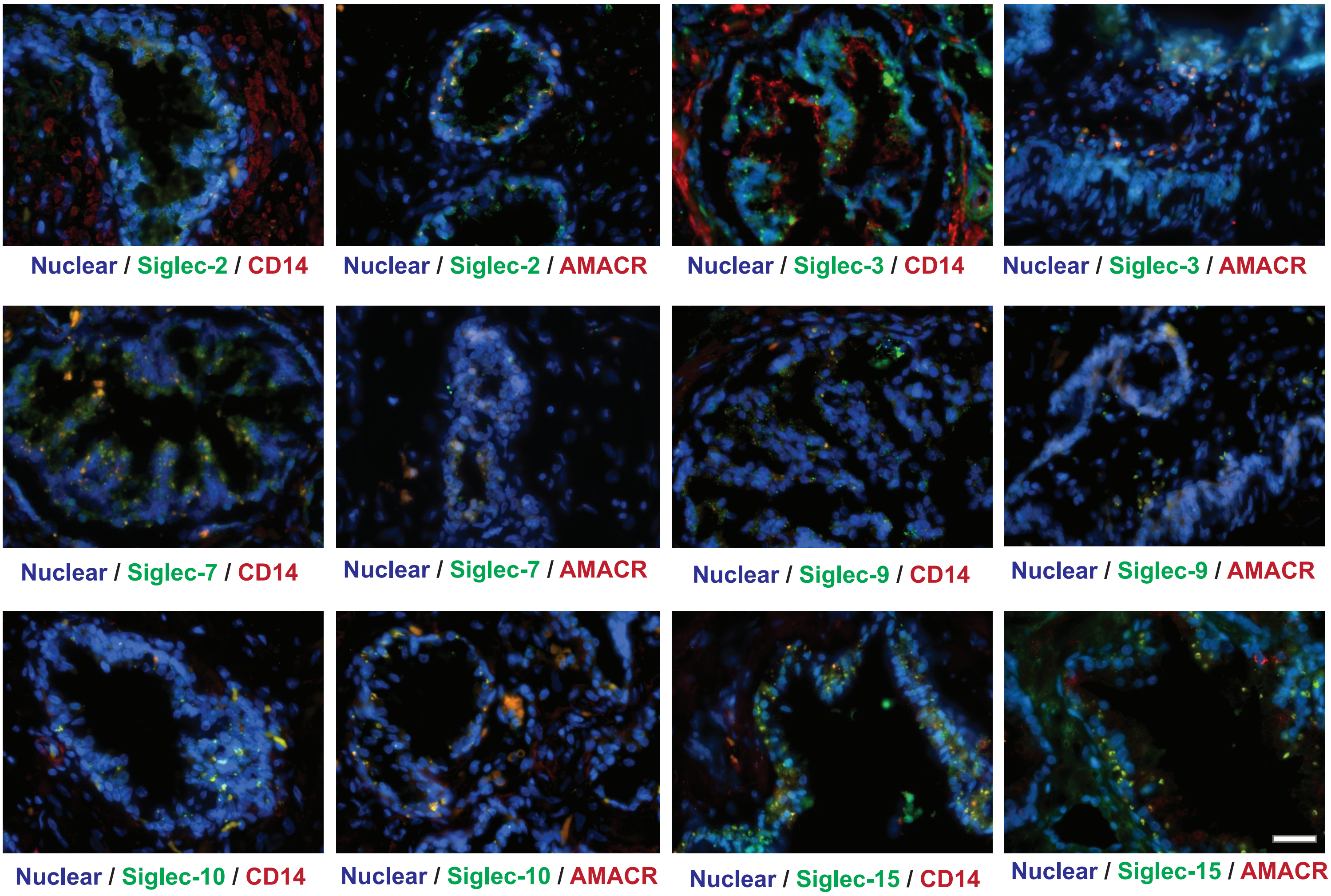

Supplementary Figure 5

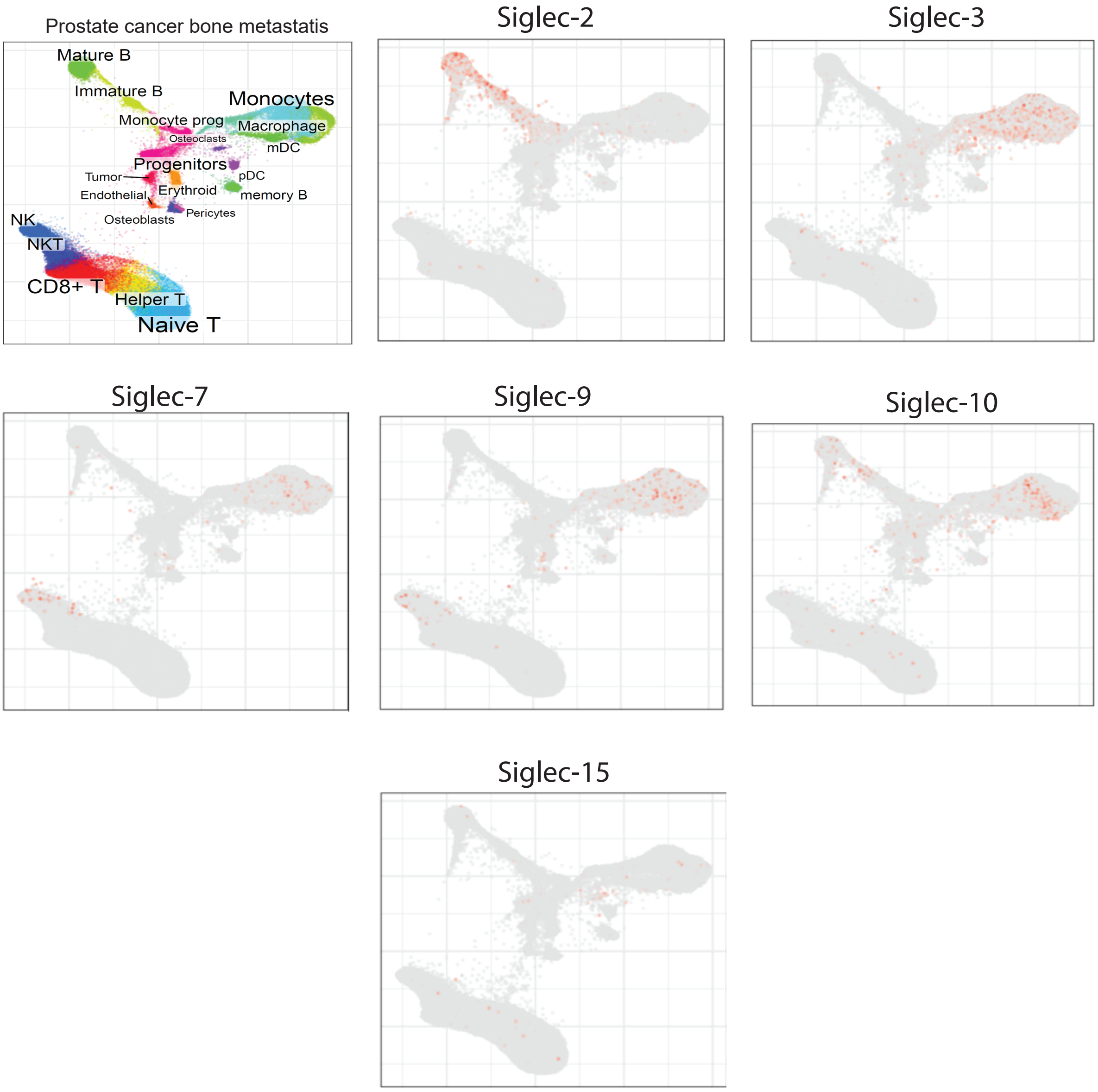

Supplementary Figure 6

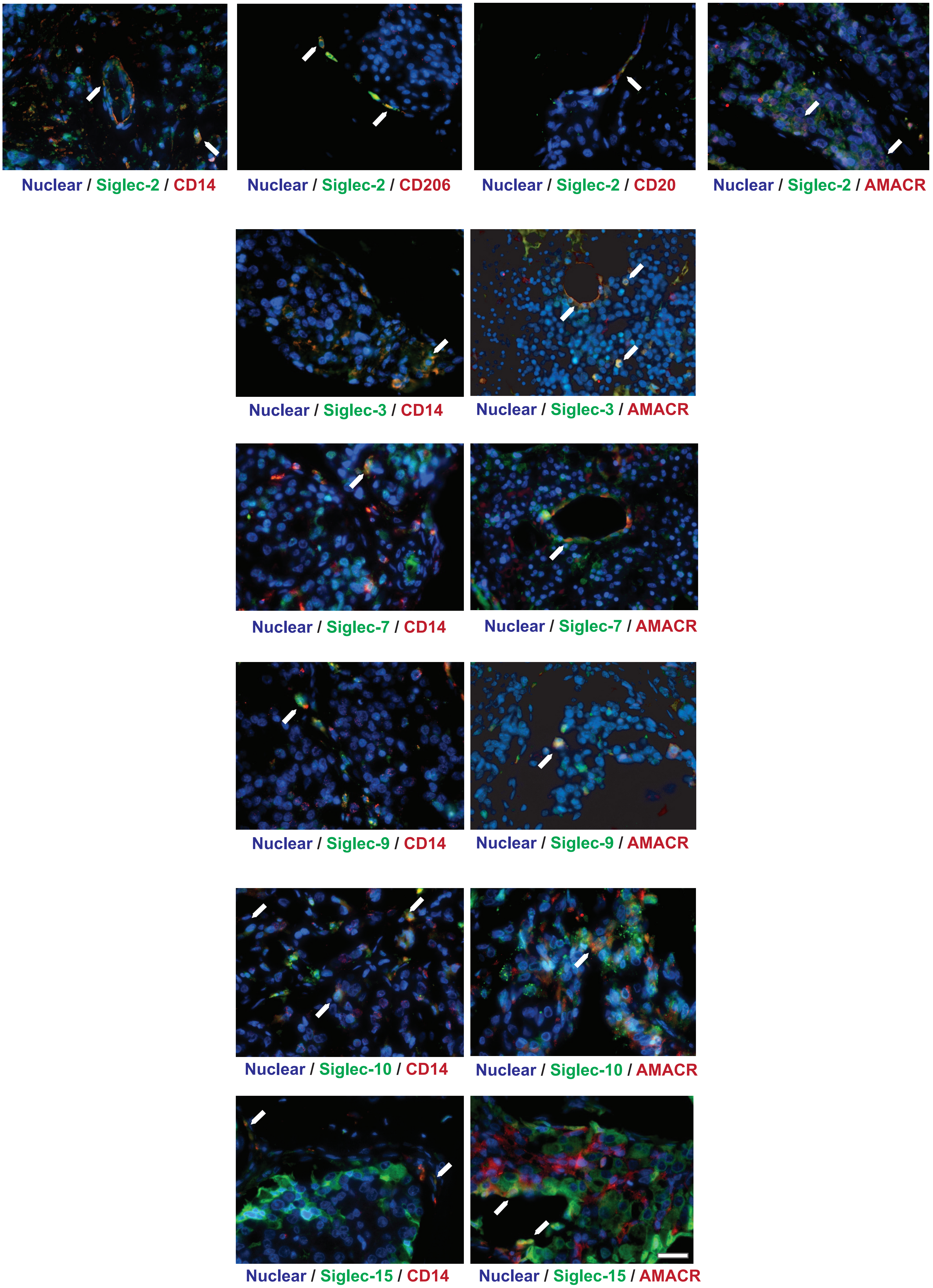

Supplementary Figure 7

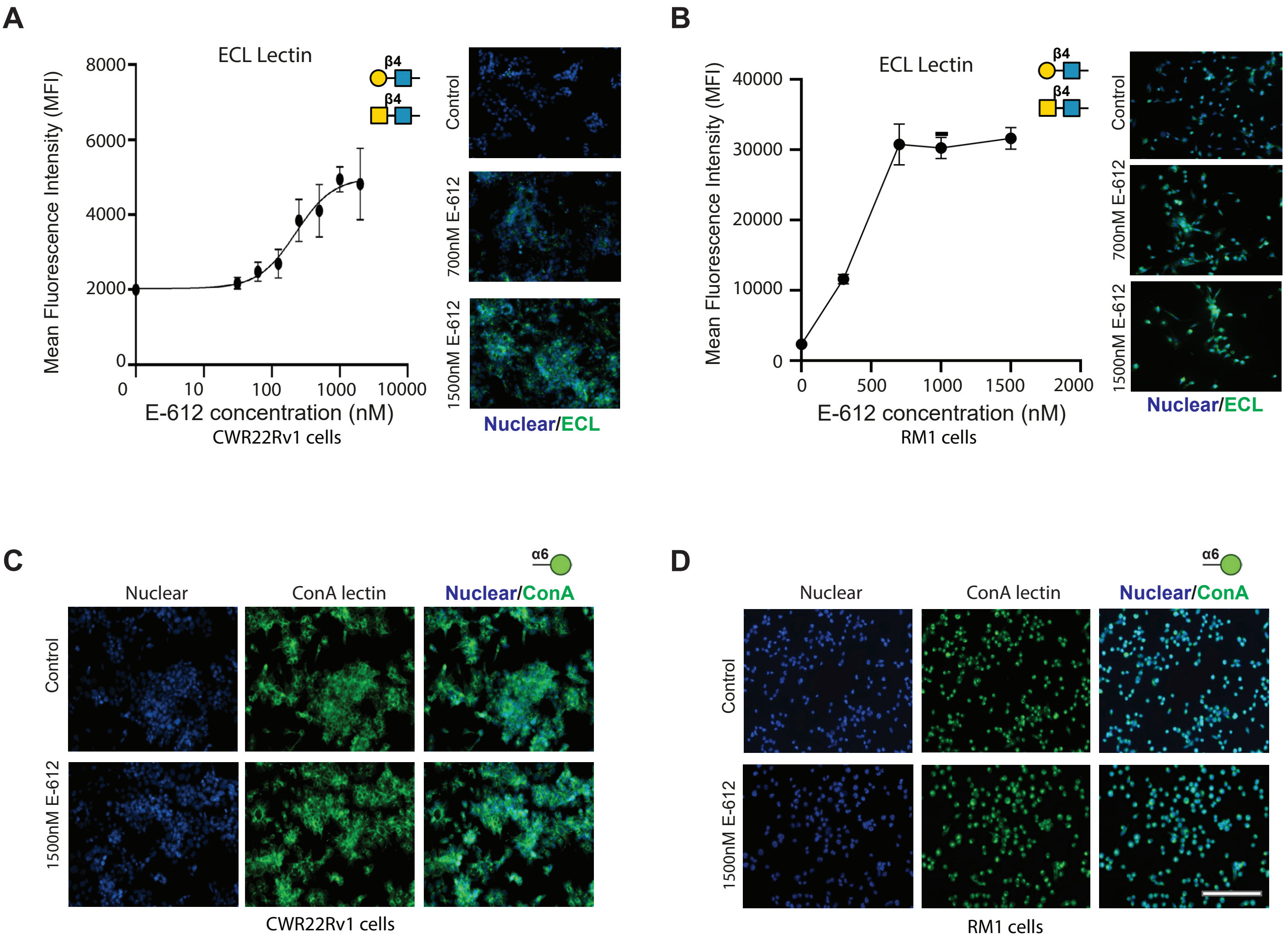
